# Deep Brain Stimulation of the Prelimbic Cortex Disrupts Consolidation of Fear Memories

**DOI:** 10.1101/537514

**Authors:** Shawn Zheng Kai Tan, Chi Him Poon, Ying-Shing Chan, Lee Wei Lim

## Abstract

Anxiety disorders pose one of the biggest threats to mental health worldwide, yet current therapeutics have been mostly ineffective due to issues with relapse, efficacy, and toxicity. Deep brain stimulation (DBS) is a promising therapy for treatment-resistant psychiatric disorders including anxiety, but very little is known about the effects of DBS on fear memories. In this study, we used a modified plus-maze discriminative task and showed that DBS of the prelimbic cortex was able to disrupt consolidation, but not acquisition or retrieval of avoidance fear memories. These results were further extended to conditioned fear memories using a standard tone-footshock fear conditioning paradigm and we demonstrated that the mechanisms were mediated by dopaminergic modulation and changes of specific neurotransmission and their metabolites in the ventral hippocampus. In conclusion, our study establishes a partial causal role of dopamine 2 receptor on the potential therapeutic role of prelimbic cortex DBS to treat anxiety disorders.

**Key Points:**

- DBS of the PrL was able to disrupt consolidation of fear memories
- We observed demonstrated short-term changes in dopaminergic receptors, c-Fos expression, and various neurotransmitters and their metabolites in the vHPC
- We established a partial causal role of dopapmine 2 receptors in the effects

## Introduction

Anxiety disorders are highly prevalent and pose one of the biggest threats to mental health worldwide (1, 2). Anxiety disorders are commonly treated using exposure therapy, a form of cognitive-behavior therapy (CBT) that targets maladaptive associated fear memories. However, such exposure therapy involves new learning that attempts to inhibit or update the previous maladaptive learning rather than erase it (3), resulting in many patients unable to maintain the benefits of CBT and often relapsing (4–9). Current attempts to enhance CBT using pharmacological or paradigm changes have met with complications such as low efficacy and drug toxicity (10, 11). Furthermore, improper administration of these techniques can lead to exacerbation of the condition (12–14).

Deep Brain Stimulation (DBS) is a surgically invasive technique that involves implanting electrodes in specific regions of the brain to modulate firing of neurons through electrical stimulation. It has been shown to be a promising non-pharmacological treatment for depression and anxiety disorders (15–18). However, little work has been done to systematically investigate its effects on fear memory. In this study, we examined the hypothesis that DBS of the ventromedial pre-frontal cortex (vmPFC), in particular the prelimbic cortex (PrL), a structure we have previously argued to be an ideal target (19), is able to disrupt fear memories. Both innate and conditioned fear were investigated in rats using a modified avoidance plus maze in which bob cat urine (aversive odour) was placed in one closed arm of an elevated plus maze and rabbit urine (neutral odour) was placed in the opposite closed arm. The results were further extended using a standard tone-footshock conditioning paradigm. We then probed the role of the hippocampus in the effects seen, and showed that ventral hippocampal dopaminergic transmission plays a central role in PrL DBS ability to disrupt consolidation of fear memories.

## Results

### DBS of PrL disrupts consolidation, but not acquisition or retrieval of avoidance fear memory

To investigate the effect of PrL DBS on avoidance fear memory, rats implanted with bilateral electrodes in the PrL were tested in a modified EPM with aversive odor in one closed arm, and then subjected to the retrieval task without odor 24h later. Rats were either stimulated for 5min in the home cage before continual stimulation in the modified EPM (Acquisition, Fig. 1A); stimulated for 5min in the home cage before continual stimulation in the EPM during the retrieval task (Retrieval, Fig. 1F); or stimulated for a further 15min at 15min after the acquisition task (Consolidation, Fig. 1K).

**Figure 1.**
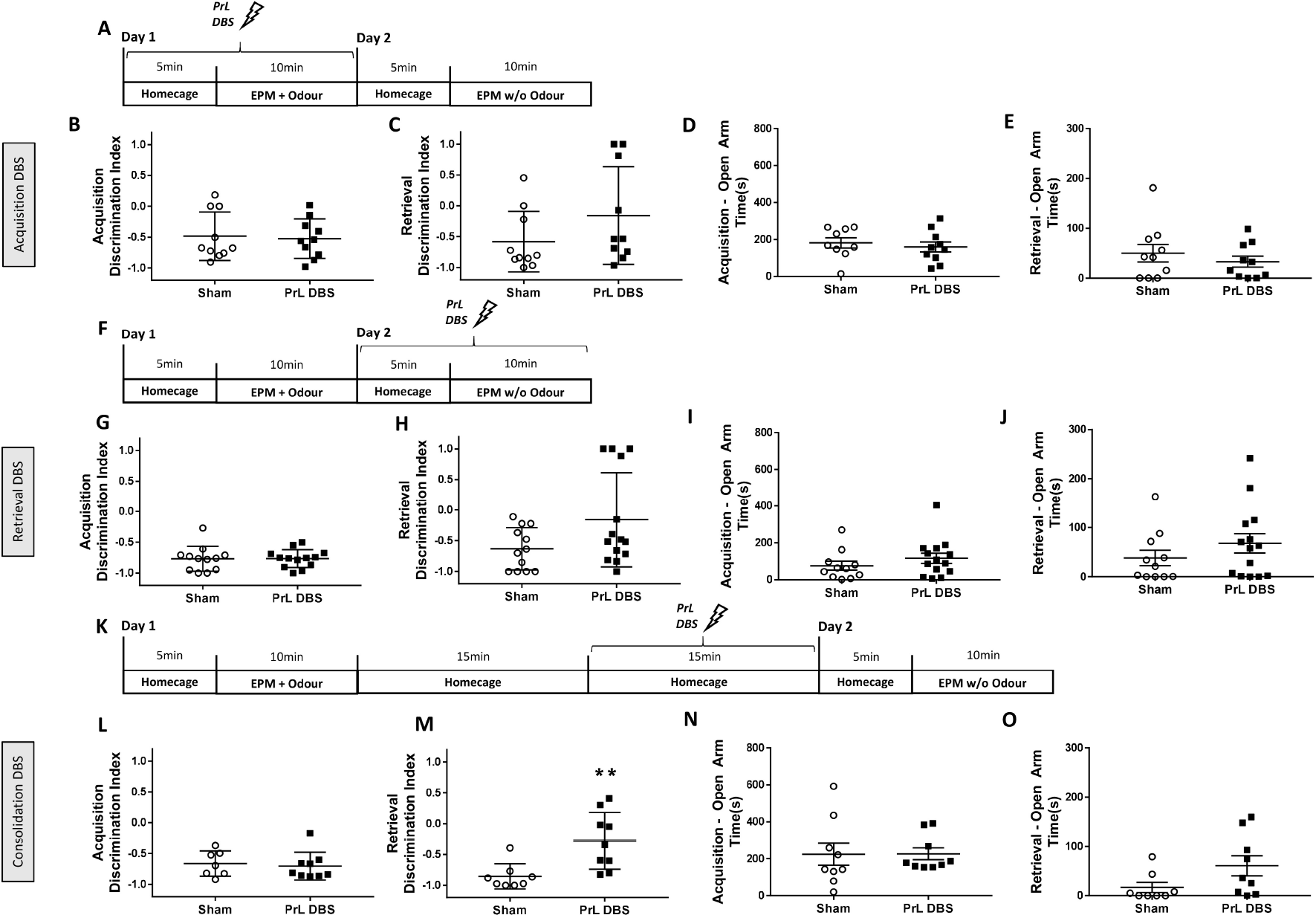
PrL DBS affects consolidation but not acquisition or retrieval of fear memory. Rats were trained in a modified elevated plus maze with aversive odor in one closed arm, and neutral odor in the opposite closed arm. Rats were tested for arm preference 24h after acquisition. Rats were stimulated either during acquisition (n=20) (A), retrieval (n=27) (F), or consolidation (n=18) (K). Both acquisition stimulation (A, B) and retrieval stimulation (G, H) showed no significant differences in the discrimination index of the closed arm with aversive odor vs neutral odor during the acquisition and retrieval tests, indicating no effect on memory. For the consolidation stimulation, L shows there was no signifreicant difference in the discrimination index during acquisition, whereas B shows significant differences in the retrieval test, indicating disruption of consolidation of memory. No significant differences were seen in the time spent in the open arms in both the acquisition and retrieval tasks for the acquisition DBS group (D, E), and retrieval DBS group (I, J). For the consolidation stimulation, N shows there was no significant difference in the time spent in the open arms, whereas O shows a significant difference in the time spent in the open arms during the retrieval task. * p<0.05, ** p<0.01.

For the acquisition DBS group (n=20), unpaired t-test of the Discrimination Index (DI) on day 1 showed no significant difference (t(18)=0.24, p=0.81), indicating that PrL DBS had no effect on acquisition of avoidance fear memory (Fig. 1B). Similarly, DI on day 2 showed no significant difference (t(18)=1.44, p=0.17), indicating that PrL DBS during acquisition had no effect on retention of avoidance fear memory (Fig. 1C). There were no significant differences in the time spent in open arm (t<0.82, all p>0.05) (Fig. 1D,E), or distance (t<0.62, all p>0.05) (Supp. Fig. 1A,B), indicating that PrL DBS did not affect innate fear or locomotion.

For the retrieval DBS group (n=27), unpaired t-tests showed no significant differences in both acquisition (t(23)=0.05, p=0.96) and retrieval (t(24)=1.97, p=0.06) tasks (Fig. 1G,H). While Fig 1G shows a p-value close to significance, this is due to the 4 animals with DIs close to 1.0, which shows overgeneralized fear rather than lack of avoidance fear memory (in which case the DI would sit closer to 0 indicating no side preference). Overall results indicate that PrL DBS has no effect on retrieval of avoidance fear memory. Similarly, no significant differences were seen in the time spent (t<1.14, all p>0.05) (Fig. 1I,J) or distance moved (t<0.95, all p>0.05) in the open arms on either day (Supp. Fig. 1D,E).

For the consolidation DBS group (n=18), unpaired t-test of the DI in acquisition task showed no significant difference (t(14)=0.37, p=0.72)(Fig. 1L). Unpaired t-test of the DI on day 2 retrieval task showed a significant difference (t(15)=3.25, p=0.005), suggesting that PrL DBS was able to disrupt consolidation of avoidance fear memory (Fig. 1M). There was no significant difference in time spent in the open arms on either day (t<1.83, all p>0.05) (Fig. 1N,O), suggesting the difference seen was not due to differences in innate fear. Unpaired t-test of the distance travelled on day 1 showed no significant difference (t(14)=0.45, p=0.66) (Supp. Fig. 1H), whereas the distance travelled on day 2 showed a significant difference (t(16)=2.17, p=0.045) (Supp. Fig. 1I). Taking into account all previous experiments, these results seem to indicate that DBS did not cause any locomotion differences, but rather PrL DBS animals had higher exploratory drive.

### Single PrL stimulation during consolidation disrupts both tone and contextual conditioned fear memory

To investigate if previous results (disrupting consolidation) can be generalized to conditioned fear memories, rats (n=18) implanted with bilateral electrodes in the PrL were subject to a standard tone-footshock fear conditioning paradigm. 15min after the acquisition task, rats were stimulated (or sham stimulated) in the home cage for 15min. 24h later, animals were placed back in the same context to test contextual fear memory. 24h after the context test, animals were subjected to a new context with five tones without footshock to test tone-footshock memory.

Two-way repeated measures ANOVA of the percentage of freezing in conditioning trial and inter-trial intervals (ITIs) revealed an effect in the CS-US trial (F(2,32)=72.1, p<0.001; F(3,48)=117.9, p<0.001), but not treatment or interaction (F<2.9, all p>0.05) (Fig. 2A,B). This indicates proper acquisition of tone and context fear memory with no difference between groups observed prior to DBS. Unpaired t-test of the context test showed a significant difference between sham and DBS groups (t(15)=2.26, p=0.04) (Fig. 2C). Unpaired t-test of the exploration period for tone test context showed no significant differences between groups (t(14)=0.22, p=0.83), indicating similar baseline levels for new context (Fig. 2E). In tone test, unpaired t-test of the average percentage of freezing for all five tones showed a significant difference between sham and DBS groups (t(13)=3.11, p=0.008) (Fig. 2D). Animals were subsequently placed in an open field 24h after the tone test to ensure the effects were not due to differences in locomotion. Unpaired t-test of distance travelled showed no significant differences between sham and DBS groups (t(17)=1.27, p=0.22), indicating no effects on locomotion (Fig. 2F).

**Figure 2.**
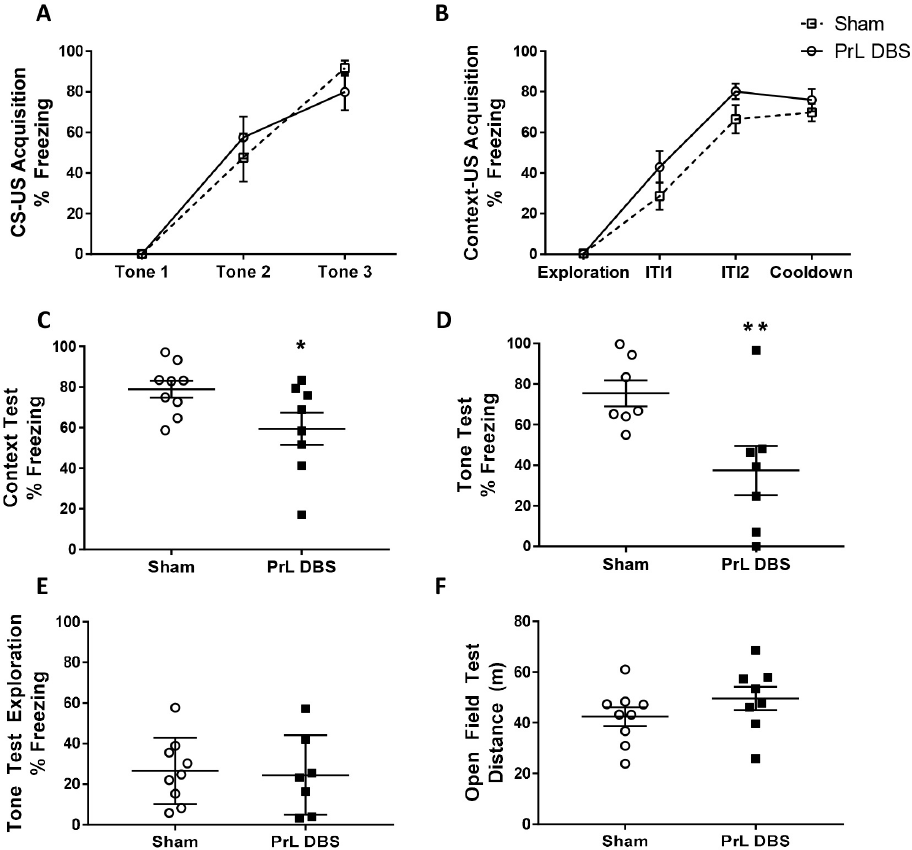
PrL DBS disrupts consolidation of both contextual and tone-footshock memory. Rats (n=19) were fear conditioned in a tone-footshock paradigm. PrL DBS was carried out 15min after acquisition. Rats were then tested for contextual fear memory and tone-footshock memory. A B shows there was effective learning of both CS-US and context-US fear memory, with no differences between groups at baseline. Figure C shows there was significantly less freezing in the context test, whereas D shows significantly less freezing in the tone test. E shows no differences in group during tone test exploration indicating that freezing was not generalized. F shows there was no significant difference in the distance travelled in the open field test, indicating no difference in locomotion. * p<0.05, ** p<0.01.

### Single stimulation during consolidation alters expressions of Drd2, Grm5, GluN2A receptor, and c-Fos, expression in the vHPC

To understand the molecular mechanisms of PrL DBS on the hippocampus, rats were sacrificed immediately after the trials on day 2 of the modified EPM. RT-qPCR was performed to detect genes related to learning and memory (23, 26, 33–35) in dHPC and vHPC sections. t-tests showed no significant differences in any of the detected genes in the dHPC and vHPC (t<1.74, all p>0.05) (Supp. Fig. 2A,B), indicating 15min of PrL DBS did not cause long-term changes in gene expressions.

To examine the immediate changes in receptor expressions in the hippocampus upon PrL DBS, rats (n=20) subjected to a similar trial as in the consolidation DBS group of the modified EPM were immediately sacrificed after DBS (Fig. 3A). RT-qPCR was performed to detect mRNA changes in dHPC and vHPC sections (Fig. 3B). t-tests of fold changes of genes in the dHPC showed no significant differences (t<1.65, all p>0.05) (Fig. 3C,D). t-tests of fold changes in the genes of the vHPC showed significant changes in the expressions of Drd2 (t(18)=2.37, p=0.029), Grm5 (t(18)=2.12, p=0.048), and GluN2A (t(18)=2.19, p=0.041), but not Drd1, Grm2, Grm3, or GluN2B (t<0.17, all p>0.05) (Fig. 3D).

**Figure 3.**
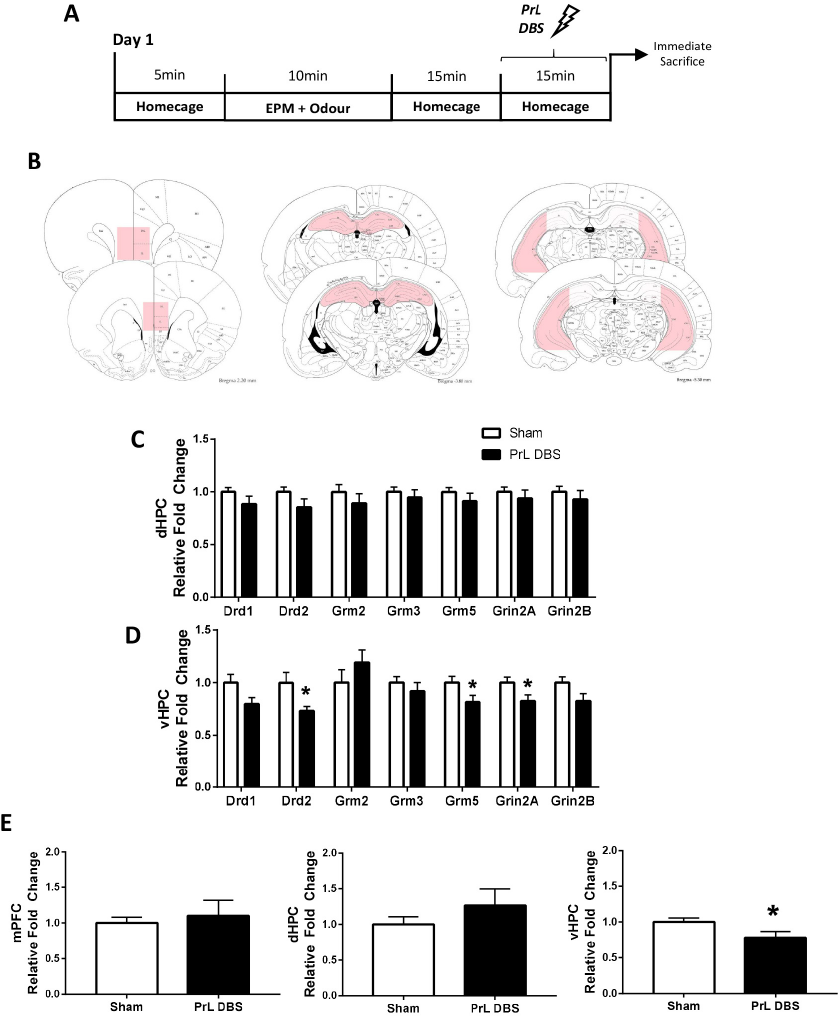
qPCR of various memory related-and neuronal activity-related genes immediately after stimulation. Rats (n=20) were trained in a modified elevated plus maze with aversive odor in one closed arm and neutral odor in the opposite closed arm. PrL DBS was carried out 15 min after acquisition and rats were immediately sacrificed (A). mPFC, dHPC, and vHPC sections were micro-dissected for qPCR (B). C shows there were no significant differences in the dHPC. D shows downregulation of Drd2, Grm5, and Grin2A in the vHPC. C-Fos gene expression was not significantly changed in the mPFC or dHPC (E and F) but was significant downregulated in the vHPC (G). * p<0.05.

To detect the changes in neuronal activity of PrL DBS in the hippocampus, the immediate early gene, c-Fos, was examined in mPFC, dHPC, and vHPC sections by RT-qPCR. t-tests of the fold changes in c-Fos expression showed no significant differences in the mPFC and dHPC (t<1.12, all p>0.05), but there was a significant decrease in the vHPC (t(17)=2.16, p=0.045) (Fig. 3E).

### vHPC dopamine 2 receptors are involved in the effects of PrL DBS on consolidation of memory

To establish the causal role of vHPC D2 receptor in the effects of PrL DBS on consolidation, rats (n=69) were implanted with electrodes in the PrL and guide cannulas in the vHPC. Rats subjected to the modified EPM were then administered either aCSF, Quinpirole (a D2R agonist), or Raclorpide (a D2R antagonist) via the guide cannula in the vHPC. At 15min after the acquisition task, rats were stimulated (or sham stimulated) in the home cage for 15min. Rats underwent the same EPM without odor 24h later to test retention of fear memory (Fig. 4A).

**Figure 4.**
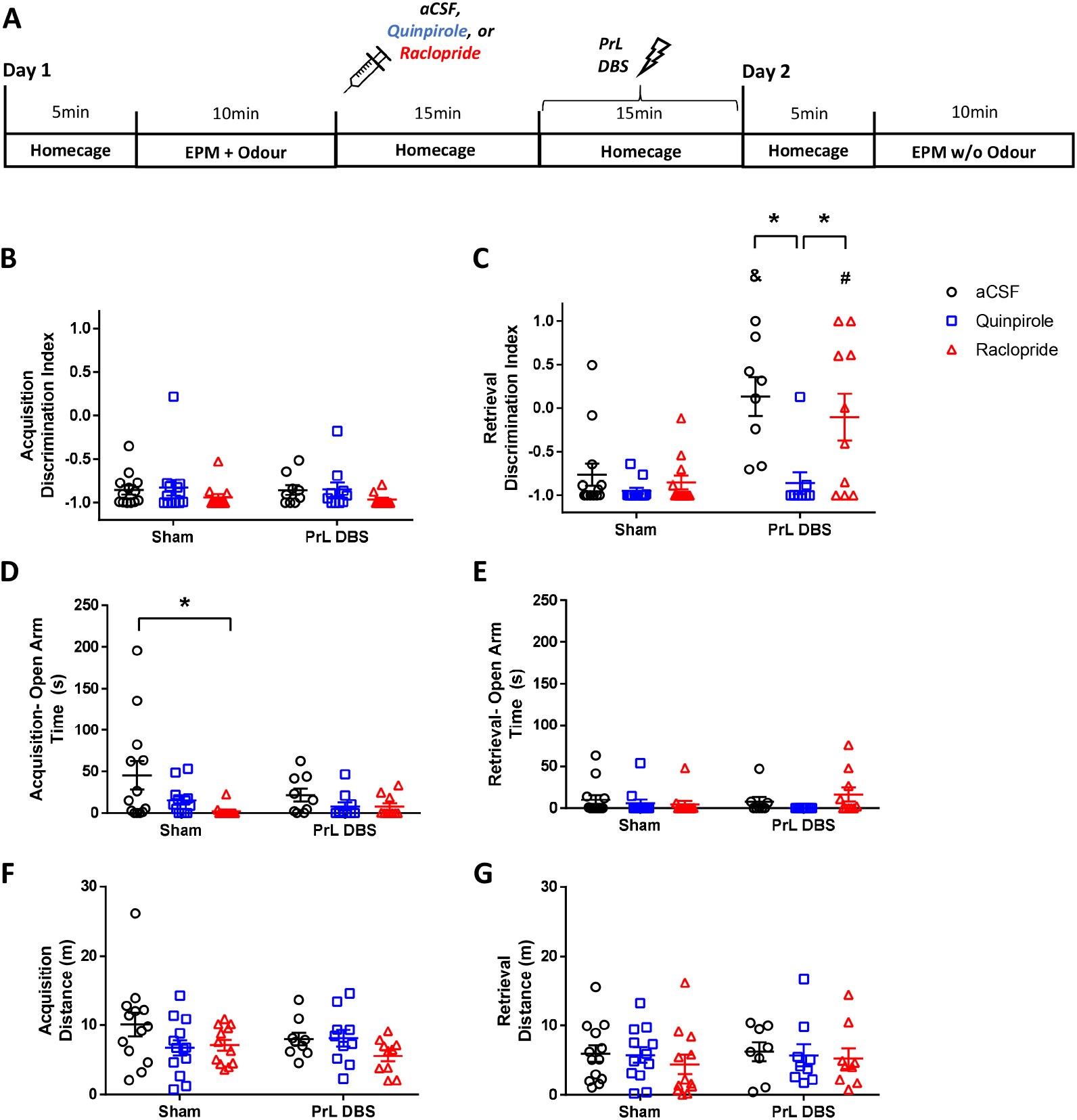
vHPC Dopamine 2 receptor is involved in the effects of DBS on consolidation of memory. Rats (n=69) were trained in a modified elevated plus maze with aversive odor in one closed arm and neutral odor in the opposite closes arm. aCSF, Quinpirole, or Raclopride were immediately administered to the vHPC. PrL DBS was carried out 15min after acquisition (A). B shows there was no significant difference in the discrimination index during acquisition, whereas C shows significant difference between DBS and sham groups for aCSF and Raclopride, but not Quinpirole group. None of the sham groups showed significant differences with each other. Quinpirole DBS group was significantly different from the aCSF and Raclorpide groups (C). There was a significant difference between the aCSF and Raclopride sham pretreatment groups (D), but no significant difference was observed in the retrieval task (E), which suggests minor batch differences, but overall no effect on innate fear. No significant difference was seen in the distance travelled indicating no effects on locomotion (F, G).

Two-way ANOVA of the DI in the acquisition task showed no significant differences (F<1.8, all p>0.05) indicating the baseline fear between the two groups were similar (Fig. 4B). Two-way ANOVA of the DI in the retrieval task showed an effect for interaction (F(2,58)=3.95, p=0.025), stimulation (F(1,58)=21.6, p<0.001), and drugs (F(2,58)=7.84, p=0.001). Tukey post-hoc test revealed significant differences in the aCSF sham group compared to aCSF DBS (p=0.002) and Raclopride DBS (p = 0.026) groups, but not Quinpirole DBS group. Within the DBS groups, aCSF and Raclopride groups showed a significant difference compared with the Quinpirole group (aCSF: p=0.001; Raclopride: p=0.016), but not with each other (p=0.91). All sham groups showed no significant differences with each other (lowest p=0.93) (Fig. 4C). Two-way ANOVA of the time spent in the open arms in the acquisition task showed an effect for drugs (F(2,61)=5.07, p=0.01), but not interaction or stimulation (F<1.26, all p>0.05). Tukey post-hoc test revealed a significant difference in only the aCSF sham group compared with Raclopride sham group (p=0.01) (Fig. 4D). However, this effect disappeared in the retrieval task, with the two-way ANOVA of the time spent in open arms showing no effects (F<1.52, all p>0.05) (Fig. 4E). The differences seen in the acquisition task could be attributed to either batch or random effects, although given the small differences in the actual mean time (around 40s) and no differences in the retrieval task, we believe the results are still valid. Lastly, no significant difference was seen in the distance travelled in both the acquisition and retrieval tasks (F<2.50, all p>0.05) (Fig. 4F,G).

### PrL DBS modulates neurotransmitters in the vHPC

To understand the effects of PrL DBS on neurotransmission, GC/MS was performed for Glu, GABA, HVA, DOPAC, DA, 5-HIAA, and 5-HT in mPFC, dHPC, and vHPC slices. All targets were within the linear ranges of the standard curves, except for DA, which was excluded from the analysis as it was only detected in the vHPC of vmPFC DBS group. t-test of the relative fold changes (target/average sham) in the vHPC sections revealed significant differences in all targets (t>3.49, p<0.05) with decreases in GABA, Glu, and 5-HIAA, and increases in HVA, DOPAC, and 5-HT (Fig. 5C). The t-test of the relative fold changes in the dHPC sections revealed a significant decrease in only 5-HT with an increase in its metabolite 5-HIAA (t>4.22, all p<0.05) (Fig. 5B). The t-test of the relative fold changes in mPFC sections revealed a significant decrease in only 5-HIAA (t(4)=3.68, p=0.021), although the difference in the mean of only 0.07 suggests no clinically relevant significance (Fig. 5A). Chromatographs are shown in supplementary materials (mPFC: Supp. Fig. 3; dHPC: Supp. Fig. 4; vHPC: Supp Fig. 5).

**Figure 5.**
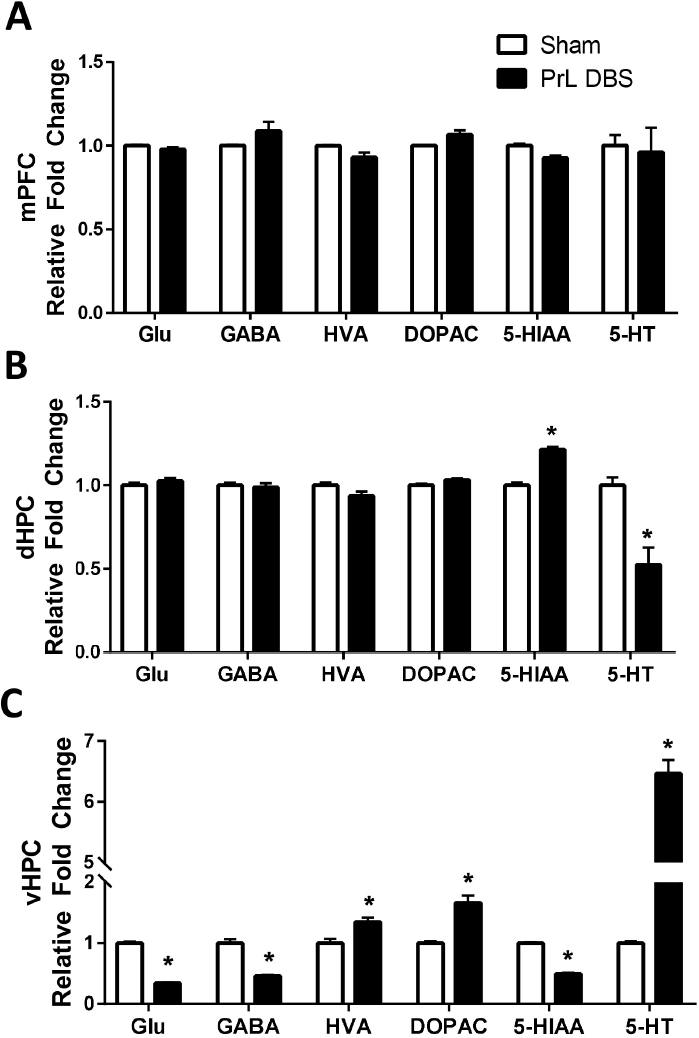
Mass spectrometry analysis of various neurotransmitters and metabolites. mPFC, dHPC, and vHPC slices were analyzed by GC/MS for Glutamate, GABA, HVA, DOPCA, 5-HIAA, and 5-HT. No Significant differences were observed in the mPFC (A), while dHPC showed significant differences with an increase in 5-HIAA and decrease in 5-HT. Significant differences were also observed in the vHPC with decreases in Glutamate, GABA, and 5-HIAA, and increases in HVA, DOPAC, and 5-HT. * p<0.05.

## Discussion

In this study, we systematically investigated the effects of acute PrL DBS on fear memory. Using our experimental parameters, we showed PrL DBS was able to disrupt consolidation, but not acquisition or retrieval of avoidance fear memory. The use of an avoidance task using a modified EPM allowed us to simultaneously control for locomotive and innate fear difference during tasks, both of which show no significant differences, suggesting an effect of memory rather than innate fear or locomotion. We further extended our results by showing similar effects in conditioned fear memory. We found changes in the expression of dopaminergic and glutamatergic receptors in the vHPC and established a partial causal role of vHPC D2 receptors in this effect. Lastly, we found changes in neurotransmitters in the vHPC.

Although using DBS to manipulate fear memory is not novel (36–39), few studies have systematically studied its effects on the individual stages of the memory process. Furthermore, most studies have focused on increasing extinction efficacy rather than directly affecting the original memory (36–38, 40). To the best of our knowledge, this is the first study to isolate the effects on the consolidation of memory by stimulating post acquisition. We also showed consistent results across several models and repeat experiments. Our findings are of great importance as the effects of neuromodulation on memories has been highly controversial, with some studies showing improvements in memory (41–44), and others showing disruption of memory (45–50).

We chose to target the vmPFC for its connections in both the hippocampus and the amygdala (51, 52), structures which are heavily implicated in fear memory (53–55). In particular, the PrL was chosen for its crucial role in learned fear (56). However, due to the proximity and functional overlap of the PrL and IL (57), a “leaked” effect to the IL cannot be entirely ruled out. We investigated the downstream effects of PrL DBS on the hippocampus and showed effects on the vHPC, but not the dHPC. This is perhaps not surprising due to the direct monosynaptic projection from the vHPC to the vmPFC, which plays a crucial role in anxiety (58) – a backpropagation of signal through DBS could be a possible mechanism. More surprising, our results suggest effects on learned rather than innate fear - the functionality of the hippocampus is thought to be split across the dorso-ventral axis, with the dHPC involved in spatial encoding and the vHPC involved in innate fear (59–63). On the other hand, the results could be related to how the dHPC and vHPC were microdissected (Fig 3B), as vHPC slices were richer in CA3 compared to dHPC slices. Obliteration of dopaminergic systems in CA3 has been shown to affect consolidation, but not acquisition of memory (64). We cannot, however, entirely exclude effects on innate fear, but rather suggest the main effect is on memory. Furthermore, the impairment of the “innate fear circuit” during consolidation might in turn lead to disruption of the consolidation of the fear aspect of the memory, which could explain the effects observed. Similarly, effects on the amygdala cannot be excluded, which probably has a role in disrupting consolidation, as indicated by Klavir et al. (65), who found that high frequency optogenetic stimulation of amygdala inputs to the PFC disrupted consolidation, but not acquisition of fear memory. Although optogenetic stimulation is mechanistically different from DBS, it shares the same concept of axonal activation (66), that could suggest another mechanism by which DBS can disrupt fear memories. Future work on the amygdala is needed to verify this.

Based on our gene expression results, we further studied the role of vHPC dopamine 2 receptor (D2R) on the effects of DBS. It has been shown vHPC D2R is involved in spatial working memory, and there was a dose-related improvement or deficit with D2 agonist Quinpirole or D2 antagonist Raclopride respectively (23). This is in line with our current results that showed a decrease in D2R gene expression in the vHPC together with a deficit in memory. To establish a causal rather than a correlative role of vHPC D2R, we infused Quinpirole and Raclopride separately into the vHPC before DBS. Quinpirole was able to reverse the effects of DBS, whereas Raclopride alone was not able to disrupt consolidation of memory. This suggests that although vHPC D2Rs play a key role in the effects of PrL DBS on consolidation of memory, it alone is not sufficient to disrupt consolidation, or fully explain the action of PrL DBS.

One possible mechanism of how DBS exerts its action is through its modulatory effects on neurotransmitters such as monoamines (67, 68) and glutamate (69–71). We investigated how PrL stimulation during consolidation could be modulated by various neurotransmitters. GC/MS results showed changes in all tested neurotransmitters in vHPC, whereas dHPC showed only changes in the serotonin system and mPFC showing no significant differences. As both glutamate and GABA levels were lowered in vHPC, a “disruption hypothesis” might explain the observed effects, in which information flow was disrupted (72). Lowered glutamate level together with downregulated c-Fos might further suggest that PrL DBS inhibits activity in the vHPC that in turn affects memory. However, the lower GABA levels may complicate this theory and further work is needed to test this hypothesis. Similar to the study by Volle et al. (73), we showed that vmPFC DBS (in our case specifically PrL) increased 5-HT levels in the vHPC. Contrary to another study (68), we showed 5-HT was lower in the dHPC, but this could be due to differences in length of the stimulation (15min in our study compared to 4h in other study). The increase in 5-HT in the vHPC was accompanied by lower 5-HIAA, which appears to be similar to the actions of a MAO-inhibitor (74) and hints at increased 5-HT availability, although this requires further study. More pertinently and consistent with our current study, we found significant changes in dopaminergic metabolites in the vHPC, but not in dHPC or mPFC. Unfortunately, dopamine levels were too low to detect reliably, making it difficult to accurately measure the dopamine turn-over. Interestingly, we found increased dopamine metabolites in the vHPC of PrL DBS groups, and dopamine was detectable only in the PrL DBS group, but not in the sham group. This initially seems to contradict our results (in that dopamine agonist reversed the effects of PrL DBS), however, it should be noted that Quinpirole has been shown to lower dopamine and DOPAC (75), which could explain the reversing effects on PrL DBS. Further studies are needed to fully understand how vmPFC DBS affects the complex play of dopamine in modulating memory processes.

To the best of our knowledge, this is the first study to show PrL DBS can disrupt consolidation of fear memories. Our results were reproducible over multiple experiments and paradigms, consistently showing the involvement of the vHPC and dopaminergic system (particularly D2 receptors). However, the effects of DBS are complex and the precise mechanisms, including the involvement of serotonergic and glutamatergic systems, are still very much unknown and need to be further explored. Overall, we provide strong evidence and a strong basis for further research into the use of neuromodulation to disrupt fear memory processes as a possible treatment for anxiety disorders.

## Methods

### Subjects

The study was approved by the Committee on the Use of Live Animals in Teaching and Research (CULATR) of The University of Hong Kong (Ref.: 4159-16). Male Sprague-Dawley rats (n=173; 7-8 weeks old at the time of surgery) were individually housed in standard cages with food and water available ad libitum. The environmental condition was maintained at temperature (21.1 °C) and humidity (60-65%) under a reversed 12/12 h light/dark cycle.

### Surgical and Deep Brain Stimulation Procedures

Surgery and DBS procedures were performed as previously described (20, 21). In brief, a construction of bilateral platinum-iridium electrodes (0.30mm Diameter, 0.031mm area) (Synergy Engineering Pte Ltd, Singapore) was implanted in the PrL (AP: +3.0 mm; ML: +/−0.6 mm; DV: −3.6 mm) based on the Paxinos & Watson Rat Brain Atlas (22). Animals that received cannulation were also bilaterally implanted with guide cannulas in the ventral hippocampus (AP: −5.3 mm; ML: +/−5.0 mm; DV: −5.6 mm). Rats were connected to cables and stimulation was performed using a digital stimulator (Model 3800 Multi-Stim: 8-Channel Stimulator; A-M Systems, Carlsborg, USA) and two stimulus isolators (Model 3820; A-M Systems). Rats were stimulated according to the experimental parameters (100 Hz, 200 µA and 100 µs pulse width) as previously described (20, 21). The sham animals were similarly tested for behaviors, but without electrical stimulation. For verification of electrode/cannula localization tip, hematoxylin-eosin (Merck, Darmstadt, Germany) was performed to examine the implantation site. Detailed information about surgical and DBS procedures are provided in the supplementary methods.

### Administration of Drugs

Rats were infused with either Quinpirole–HCl (10 µg/side of salt, equivalent to 39.09 µmoles/side), Raclopride (1.67 µg/side of salt, equivalent 3.36 µmoles/side) or artificial cerebral spinal fluid (aCSF) at dosages previously shown to be effective in the vHPC (23). Drugs were infused into the vHPC through two Hamilton syringes (10 µl) connected to an internal cannula via polyethylene tubing (Protech International, Texas, USA). The infusion volume (2 µl) was delivered over approximately 3min and the internal cannula was left in for an additional 3min.

### Modified Elevated Plus Maze

On day 1, a container with 5 ml of Bobcat urine (aversive odor; PredatorPee, Maine, USA) was placed in one closed arm of the modified elevated plus maze (EPM) and a container with 5 ml of rabbit urine (neutral odor) was placed in the opposite closed arm. Urine was changed every 4 5 animals and testings were counter-balanced to ensure a proper dispersion of odor across tests and subjects. On day 2, empty containers without odor were placed in the two closed arms. For each trial, the animal was placed in the central platform and tested for 10min. Stimulation or sham stimulation were administered accordingly (see protocols). Discrimination Index (DI) was calculated from time spent in arm as (aversive – non aversive)/(aversive + non aversive). More details can be found in the supplementary methods.

### Real-time PCR

Immediately after experiments, animals were sacrificed, and brains were removed. The dorsal hippocampus (dHPC), ventral hippocampus (vHPC), and medial prefrontal cortex (mPFC) were micro-dissected (300 µm thickness) on a cryostat CM3050 (Leica Microsystems, Wetzler, Germany). Total RNA was extracted, reverse transcribed, and real-time PCR was performed. All primers used were previously published (24–29), and amplification efficiency was reassessed as previously described (30). Relative gene expression analysis was performed using the 2-∆∆CT method, and DBS animals were normalized to the sham animals, as previously described (30, 31). The list of primer sequences and the details of real-time PCR experiments can be found in the Supplementary Table 1 and supplementary methods.

### Mass Spectrometry

Metabolites were extracted from mPFC, dHPC, and vHPC slices, and derivatized using a standard protocol. Gas chromatography–mass spectrometry (GC/MS) analysis was performed on an Agilent 7890B GC - Agilent 7010 Triple Quadrapole Mass Spectrometer system (Agilent, California, USA). Dopamine (DA), Serotonin (5-HT), -Aminobutyric acid (GABA), Glutamic acid (Glu), 3,4-Dihydroxyphenylacetic acid (DOPAC), Homovanillic acid (HVA), and 5-Hydroxyindole acetic acid (5-HIAA) were measured with norvaline as the internal standard. More details can be found in the supplementary methods.

### Fear Conditioning

Fear conditioning was performed using a startle and fear conditioning system (Panlab Harvard Apparatus, Massachusetts, USA). For acquisition, the conditioned stimulus (CS) was a 10s tone (Volume: 80 dB, Frequency: 5000 Hz) which co-terminated with a 1s 0.6 mA footshock as the unconditioned stimulus (US). The protocol consisted of 2min of adaptation, followed by three tone-footshock pairings, and then 2min of rest before being removed from the chamber. For the context test, 24h after conditioning, rats were placed in the chamber for 5min. Percentage freezing was reported based on the 5min test duration. For the tone test, 24h after the context test, rats were placed in the chamber and tested in a different context to the one received during conditioning. The tone tests consisted of 2min of adaptation, followed by five tone presentations (10s) with 10s inter-trial interval (ITI) in the absence of footshock, and 2min of rest before being removed from the chamber. Percentage freezing was reported as the average of the five CS presentations. To assess fear learning and memory, freezing was used as the dependent variable, which is a species-specific defense response defined as the absence of all movement except that required for respiration (32). Freezing values were calculated using a high sensitivity Weight Transducer System (StartFear System, Harvard Apparatus, Holliston, Massachusetts, USA). Open Field Test was conducted 24h after the tone test to control for locomotion differences. Animals were allowed to explore the arena for 10min. The behavior of rats was recorded and analyzed using a digital video camera and Any-maze video tracking system 5.0 (Stoelting Co).

### Statistics

All statistical analyses were performed using GraphPad Prism 7.00. Statistical models used are stated in the various results sections. Outliers were removed using the ROUT method. Results were considered significant for p<0.05.

## Acknowledgements & Disclosures

The scientific work was funded by grants from the Hong Kong Research Grant Council (RGC-ECS 27104616), and The University of Hong Kong URC Supplementary Funding (102009728) that was awarded to LWL. We would like to thank the Proteomics and Metabolomics Core Facility (HKU), the Laboratory Animal Unit (HKU), as well as Anna Tse, Junhao Koh, and Smaranda Badea for their valuable input. All authors declared no biomedical financial interests or potential conflicts of interests.

