## Supplementary Methods and Materials for "Deep Brain Stimulation of the Prelimbic Cortex Disrupts Consolidation of Fear Memories"

*Surgical Procedures*

Rats were implanted with Deep Brain Stimulation (DBS) electrodes bilaterally in the Prelimbic region (PrL) of the ventromedial prefrontal cortex (vmPFC). The animals were initially anesthetized with 5% isoflurane vapor mixed with oxygen until loss of righting reflex, and were then mounted in a stereotaxic frame (Leica Biosystems, Nussloch, Germany) and maintained with 2.5% isoflurane delivered through a nose cone. A midline incision was made to expose the skull and sagittal suture was used to align the skull along the anterior-posterior axis in the frame. Paxinos & Watson Rat Brain Atlas was used as a guide for stereotaxic implantation of electrodes (from bregma: +3.0 mm anteriorposterior; +/-0.6 mm mediolateral; and -3.6 mm dorsoventral). The electrode construct was anchored to the rat skull with stainless steel screws and dental acrylic (Paladur, Heraeus Kulzer GmbH, Hanau, Germany). Animals with cannulation were also bilaterally implanted with guide cannulas in the ventral hippocampus (from bregma: - 5.3 mm anteriorposterior; +/-5.0 mm mediolateral; and -5.6 mm dorsoventral), and similarly secured with dental acrylic.

*Modified Elevated Plus Maze*

The elevated plus maze was a four arm maze made of black Plexiglass. The maze consisted of two opposing open arms (50x10 cm), and two opposing closed arms (50x10 cm) with 15 cm high walls with arms extending from the central platform (10x10 cm). On day 1, a container with 5 ml of Bobcat urine (PredatorPee, Maine, USA) as aversive odor was placed in one closed arm and a container with 5 ml of rabbit urine (neutral odor) was placed in the opposite closed arm. On day 2, containers without odors were placed in the closed arms. Rats were stimulated (or sham stimulated) accordingly for each experiment (see protocol sections). Rats were placed in the central platform and individually tested for 10 min. The time spent in the open arms, aversive closed arm, or non-aversive closed arms were recorded. The total distance moved was also measured as a marker of locomotion changes. Discrimination Index (DI) calculated as (aversive – non aversive)/(aversive + non aversive) was used as a measure of avoidance fear memory. Their behavior was recorded and analyzed using a digital video camera and Anymaze 5.0.

*Real-time PCR*

Immediately after the experiments, rats were sacrificed, and their brains were extracted and frozen in liquid nitrogen. The dorsal hippocampus (dHPC) (Bregma -3.14 mm to -3.80 mm; 4 X 100 μm) and ventral hippocampus (vHPC) (Bregma -4.80 mm to -5.30 mm; 2 X 100 μm) were dissected out in a cryostat (Leica CM3050S, Nussloch GmbH, Germany) according to the anatomical regions based on the Paxinos & Watson Rat Brain Atlas. Sections were stored at -80°C until use. Total RNA was extracted using TRIzol reagent (Molecular Research Center Inc., Ohio, USA) And then reverse transcribed using PrimeScript™ RT reagent kit with gDNA eraser (Takara Bio USA, California, USA) and cDNA products were stored at -20°C until use.

Real-time PCR was performed on a StepOne™ Real-Time PCR System (ThermoFisher Scientific, Massachusetts, USA). Reactions were performed in triplicate in MicroAmp

96-well plates with standard conditions: (50°C for 2 min, 95°C for 10 min, 40 cycles of 95°C for 10 s, 60°C for 30 s) with SYBR Green fluorescence (Applied Biosystems, Life Technologies, Warrinton, UK) was detected after each cycle. A melt curve from 60-95°C with a step increase of +5°C was plotted at the end of the cycling stage to evaluate the amplification products. Data were analyzed using StepOne™ Real-Time PCR software. The qPCR primers used were previously validated (sequences of primers and references are listed in the Supp. Table 1), and amplification efficiency was assessed for all primers as previously described (1). Hypoxanthine phosphoribosyltransferase 1 (HPRT1) was used as the internal control. Relative gene expression was calculated using the $2^{-\Delta\Delta CT}$ method, and DBS animals were normalized to the sham animals, as previously described (1, 2)

*Fear Conditioning*

Fear conditioning was conducted in a startle and fear conditioning system (Panlab, Harvard Apparatus, Massachusetts, USA). The floor of the experimental chamber (LE116; 250 mm(width) x 250 mm (depth) x 250 mm (height) consisted of a stainless steel grid spaced 10 mm apart. For the acquisition and context texts, the chamber consisted of black walls and a grid floor. A constant-current shock generator was used to deliver an electric shock to the floor of the chamber as required. For the tone test, the chamber consisted of white walls and a stainless steel plate floor. A tone generator, speaker, and sound calibration package was used to deliver auditory cues as required. Chambers were enclosed in sound-attenuating boxes in a dedicated experimental room. For acquisition, the conditioned stimulus (CS) was a tone (Volume: 80 dB, Frequency: 5000 Hz) and the unconditioned stimulus (US) was a 0.6 mA footshock. All presentations of CS and US were controlled and recorded by a computer. Rats were placed in the chamber for a 2 min adaptation period, followed by three tone-footshock pairings, and then a 2 min rest period before being removed from the chamber. Each pairing consisted of a 10 s tone co-terminated with a 1 s shock at the end. Inter-trial intervals (ITIs) were 85 s or 135 s to avoid any time associations. Freezing was calculated from the first 9 s of each CS presentation to avoid confounding effects of the shock presentation in the last second.

For the context test, 24 h after conditioning, rats were placed in the same chamber for 5 min. The percentage freezing was reported based on the entire 5 min test.

For the tone test, 24 h after the context test, rats were placed in the chamber and tested in a different context to the one received during conditioning. The tone test consisted of a 2 min adaptation period, followed by five presentations of a tone (10s tones with 10 s Inter Trial Intervals) without a footshock, and then 2 min rest period before being removed from the chamber. Percentage freezing was reported as the average of all five CS presentations.

To assess fear learning and memory, freezing was used as the dependent variable, which is a species-specific defense response defined as the absence of all movement except that required for respiration (26). Freezing values were calculated using a high sensitivity Weight Transducer System (StartFear System, Harvard Apparatus). Freezing was initially scored manually to determine the motion threshold for use in the Packwin software. All data reported in the study were automatically scored by the Packwin software with a motion threshold of 7, software gain set at 16, and hardware gain of the LE111 Load Cell Control Unit (Harvard Apparatus) set at 500.

The Open Field Test was conducted 24 h after the tone test to measure any locomotion differences. The Open Field Test consisted of a square arena (40x40 cm) with 40 cm high walls. Animals were allowed to explore the arena for 10 min And their behavior was recorded and analyzed using a digital video camera and Anymaze 5.0.

*Mass Spectrometry*

Tissue homogenization and metabolite extraction was performed in 1.5 ml of methanol/MiliQ water (80%, v/v) with 0.1 mg norvaline as the internal standard. Tissue was homogenized on ice by 10 cycles of sonication at 10 microns for 20 s and 10 s pause time. Next, 750 μl of MiliQ water was added and the tube was vortexed for 30 s, and then 1200 μl of chloroform was added and vortexed again. After agitation for15 min, the sample was centrifuged for 5 min at 10000 g. The polar phase was isolated and the dried residue was redissolved and derivatized for 2 h at 37°C in 40 μl of methoxylamine hydrochloride (30 mg/ml in pyridine), followed by trimethylsilylation for 1 h at 37°C in 70 μl MSTFA with 1% TMCS. A sample (0.2 μl) was analyzed by GC-MS and the remaining sample dried under vacuum.

The GC/MS spectra were acquired in SCAN and MRM mode on an Agilent 7890B GC - Agilent 7010 Triple Quadrapole Mass Spectrometer system (Agilent, CA, USA). The sample was separated in an Agilent DB-5MS capillary column (30 m × 0.25 mm ID, 0.25 μm film thickness) with a constant flow rate of 1 ml/min. The GC oven program started at 60°C (holding time 1 min) and increased at 10°C/min to 120°C, then 3°C/min to 150°C, followed by 10°C/min to 200°C and finally 30°C/min to 280°C (hold 5 min). Inlet temperature and transfer line temperature were 250°C and 280°C, respectively.

Characteristic quantifier and qualifier transitions were monitored in MRM mode during the run. Mass spectra from m/z 50-500 were acquired in SCAN mode.

Data analysis was performed using the Agilent MassHunter Workstation Quantitative Analysis Software. Linear calibration curves for each analyte were generated by plotting the peak area ratio of external/internal standard against the standard concentration at different concentration levels. Analytes were confirmed by comparing the retention time and ratio of characteristic transitions between the sample and standard.

*Statistical Analysis*

All statistical analyses were performed using GraphPad Prism 7.00. Statistical models used are given in the appropriate results sections. Outliers were removed using the ROUT method (1%). Results were considered significant for $p<0.05$.

### Supplementary Table

| Target Gene | Sense | Anti-Sense | References |
|---|---|---|---|
| **Drd1** | 5'-CCTTCGATGTGTTTGTGTGG-3' | 5'-GGGCAGAGTCTGTAGCATCC-3' | Dick et., 2015 (3) |
| **Drd2** | 5'-TTCTGTCCTTCACCATCTCC-3' | 5'-GACCAGCAGAGTGACGATGA-3' | Dick et al., 2015 (3) |
| **Grm2** | 5'-AGTCCTTAGCTGGGGAGCCT-3' | 5'-AACCATCCTCTCTATCCCAGAGTAAC-3' | Ermolinsky et al., 2008 (4) |
| **Grm3** | 5'-TAGGCTGTTAGACAAAGTGCTCA-3' | 5'-GAAGGGGCTGTTAATTAGGGCA-3' | Ermolinsky et al., 2008 (4) |
| **Grm5** | 5'-ACCAAGACCAACCGTATTGC-3' | 5'-AGACTTCTCGGATGCTTGGA-3' | Tan et al., 2015 (5) |
| **Grin2a** | 5'-GCACCAGTACATGACCAGATTC-3' | 5'-ACCAGTTTACAGCCTTCATCC-3' | Calabrese et al., 2012 (6) |
| **Grin2b** | 5'-TTCATGGGTGTCTGTTCTGG-3' | 5'-GGATGTTGGAGTGGGTGTTG-3' | Calabrese et al., 2012 (6) |
| **c-Fos** | 5'-CCGACTCCTTCTCCAGCAT-3' | 5'-TCACCGTGGGGATAAAGTTG-3' | Rogers et al., 2004 (7) |
| **HPRT** | 5'-CTCATCGGACTGATTATGGACAGGAC-3' | 5'-GCAGGTCAGCAAAGAACTTATAGCC-3' | Covacu et al., 2009 (8) |

**Supplementary Table 1. List of primers used in qPCR.** Table of primer sequences for qPCR with references. All primers were tested for efficiency before use.

**Supplementary Figures**

**Supplementary Figure 1. PrL DBS does not affect locomotion.** No significant differences were seen in the distance travelled for both the acquisition and retrieval tasks in the acquisition DBS group (A, B) and retrieval DBS group (D, E). For consolidation stimulation, H shows there was no significant difference in the distance travelled, whereas I shows a significant difference in distance travelled, suggesting higher exploratory drive rather than a difference in locomotion. C, F, & J show electrode implantation sites. * $p<0.05$.

**Supplementary Figure 2. qPCR of various synaptic plasticity- and neuronal activity-related genes 24h after stimulation.** Rats from the consolidation experiment (Fig. 1K) were sacrificed, and dHPC and vHPC sections were micro-dissected for qPCR. No significant differences were seen in the detected gene expressions in both dHPC and vHPC (A, B).

**Supplementary Figure 3. Mass spectrometry chromatographs of various neurotransmitters and metabolites in the mPFC.** mPFC slices were analyzed by GC/MS for Glutamate, GABA, HVA, DOPCA, 5-HIAA, and 5-HT. A shows chromatographs of Sham group, while B shows chromatographs of PrL DBS group.

**Supplementary Figure 4. Mass spectrometry chromatographs of various neurotransmitters and metabolites in the dHPC.** dHPC slices were analyzed by GC/MS for Glutamate, GABA, HVA, DOPCA, 5-HIAA, and 5-HT. A shows chromatographs of Sham group, while B shows chromatographs of PrL DBS group.

**Supplementary Figure 5. Mass spectrometry chromatographs of various neurotransmitters and metabolites in the vHPC.** vHPC slices were analyzed by GC/MS for Glutamate, GABA, HVA, DOPCA, 5-HIAA, and 5-HT. A shows chromatographs of Sham group, while B shows chromatographs of PrL DBS group.
