## Supplementary figures and images for "Deep Brain Stimulation of the Prelimbic Cortex Disrupts Consolidation of Fear Memories"

### Supp Fig 1

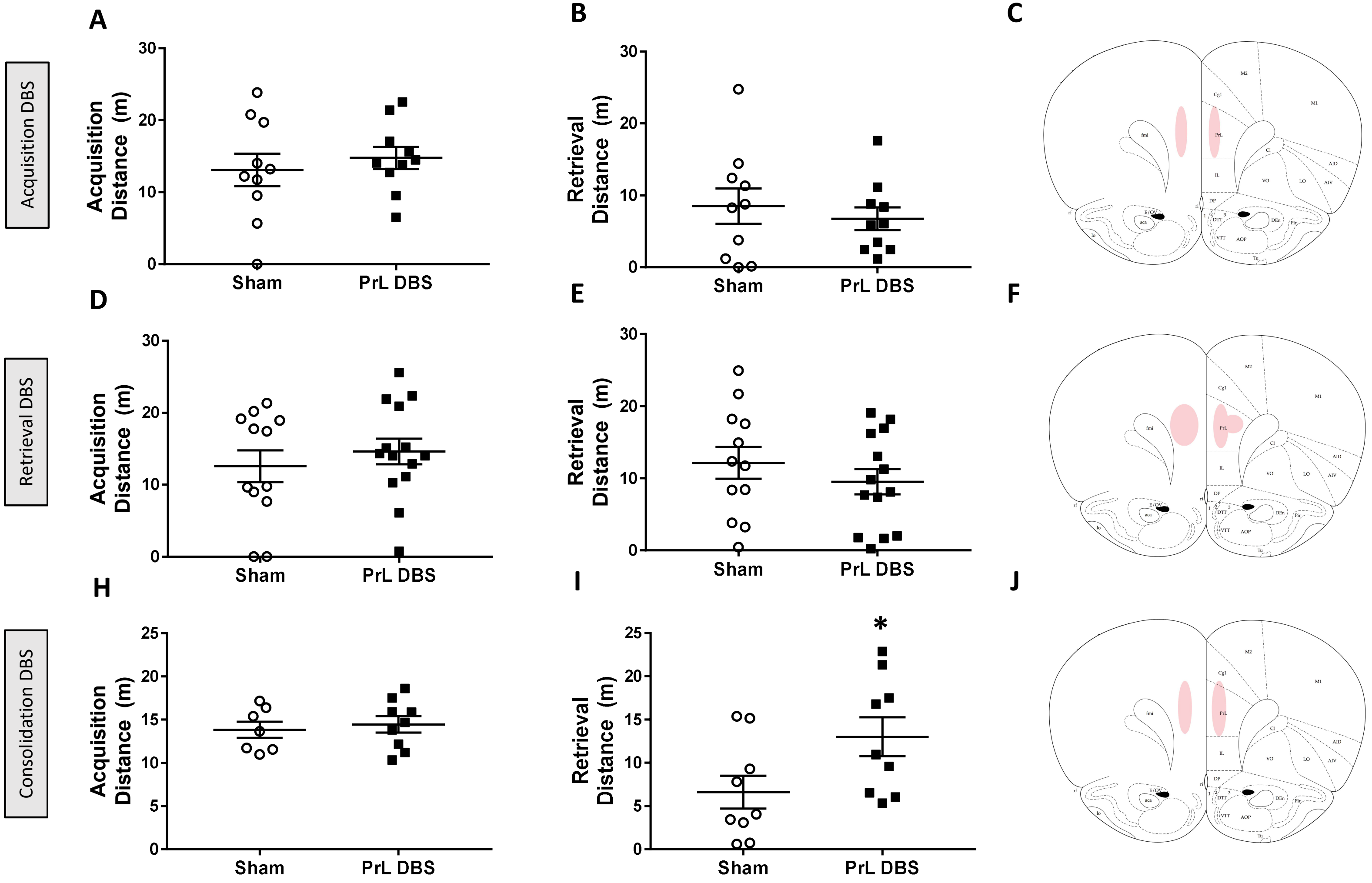

### Supp Fig 2

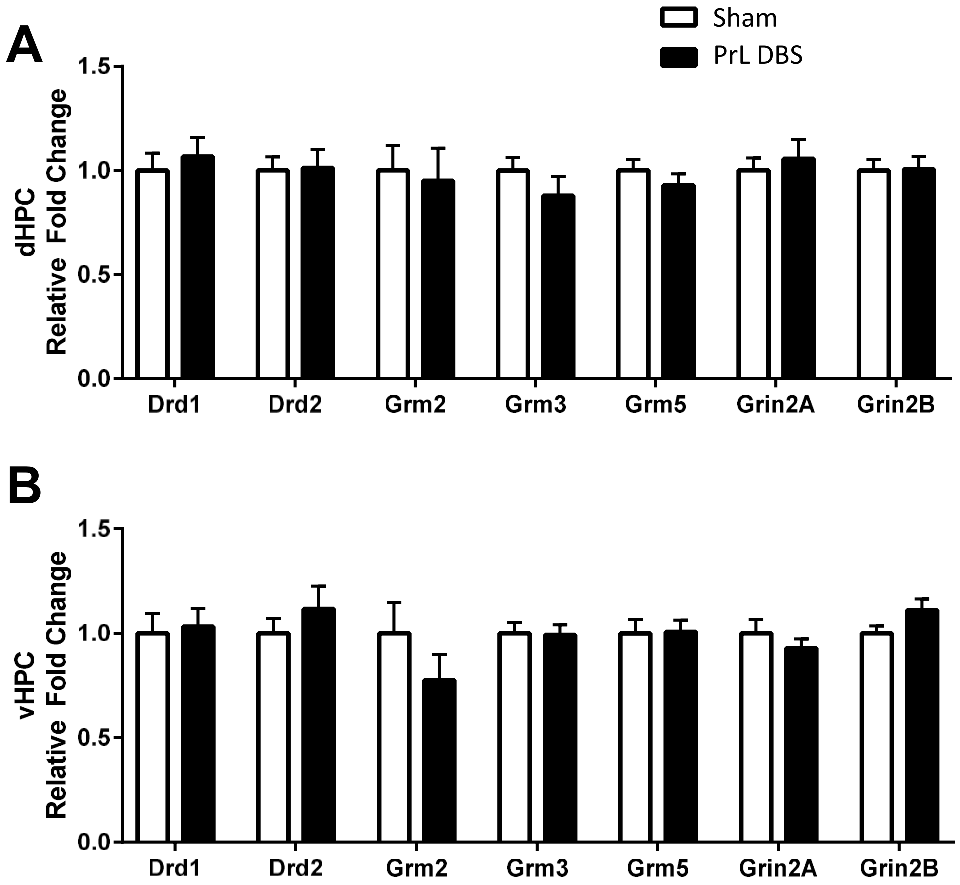

### Supp Fig 3

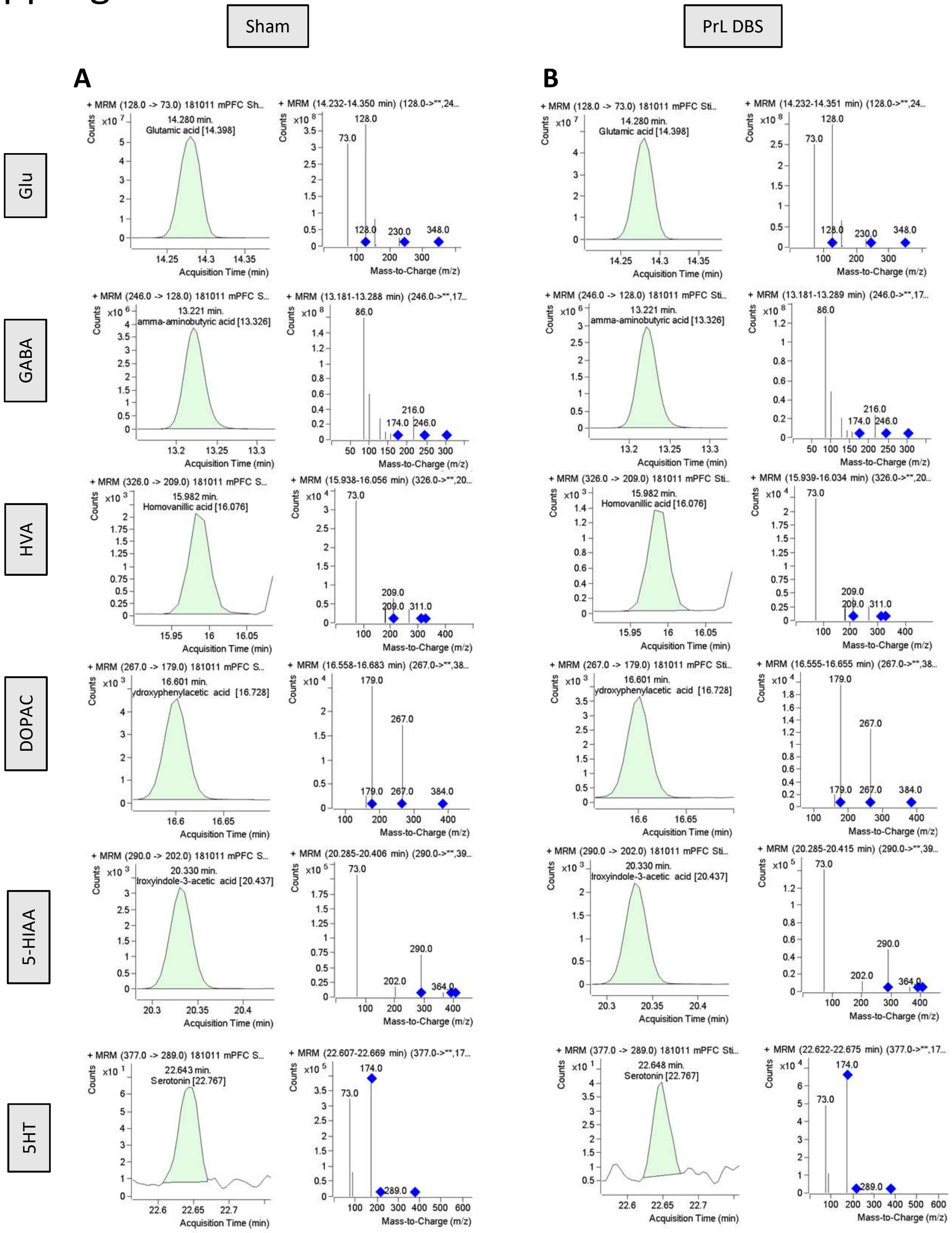

### Supp Fig 4

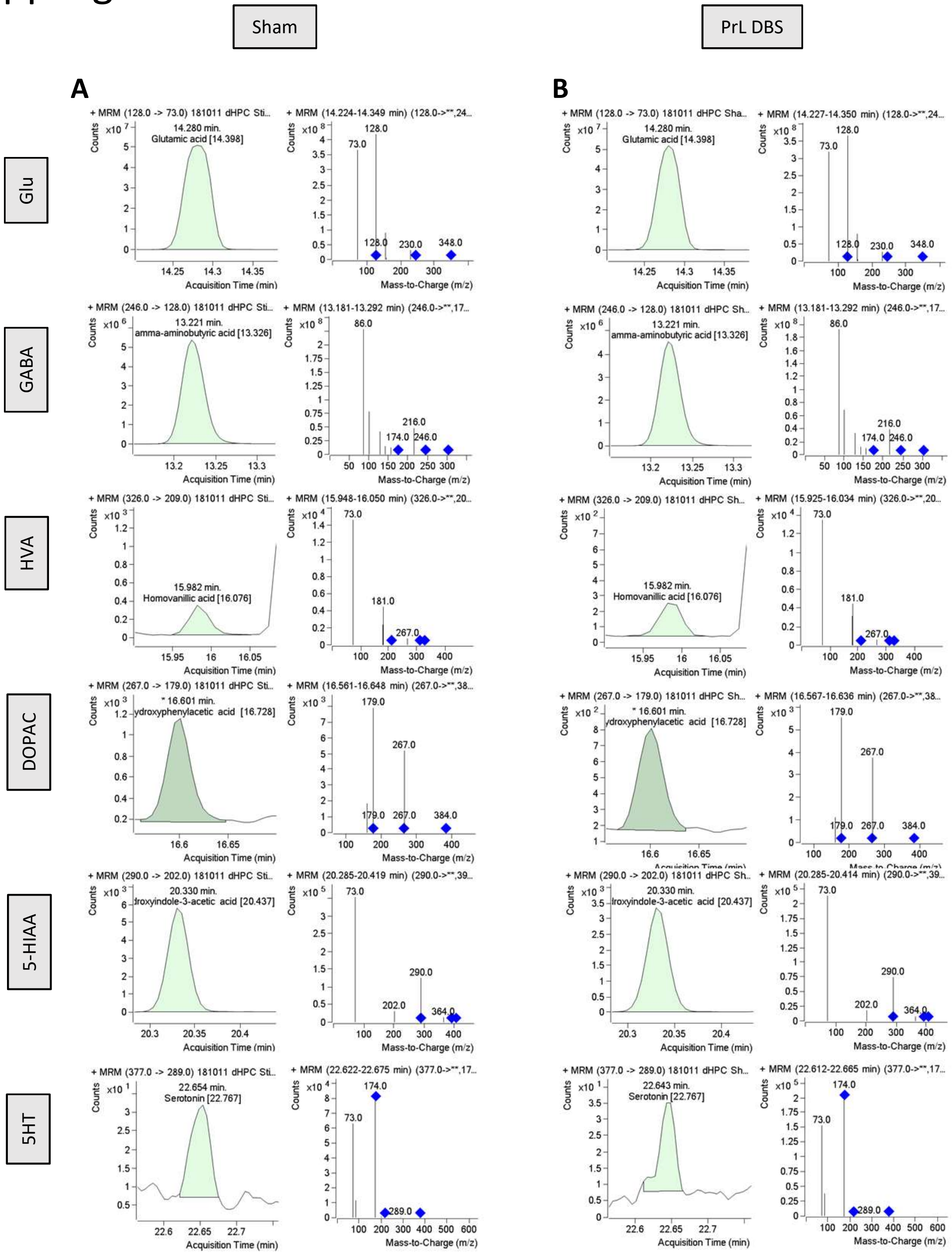

### Supp Fig 5

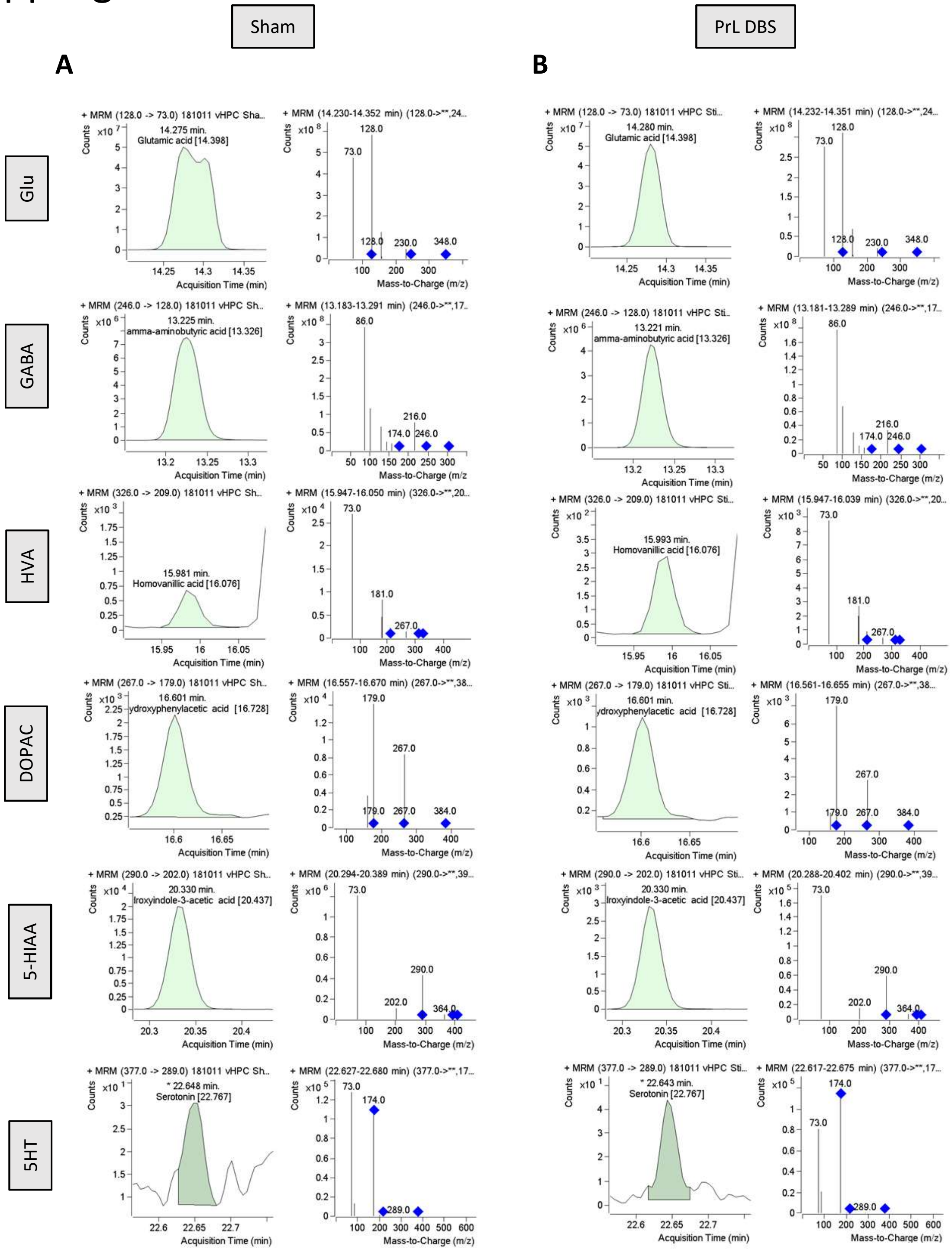
